# Time-Series RNA Sequencing Reveals Temperature-Specific and Temperature-Compensated Circadian Genes in *Drosophila*

**DOI:** 10.1101/2025.11.04.686609

**Authors:** Fumihiro Ito, Fei Zhang, Wanhe Li

**Author notes:** Corresponding author: Wanhe Li.

## Abstract

Circadian rhythms align the physiology and behavior of living organisms with the 24-hour day. As a defining property of circadian rhythms, temperature compensation preserves the 24-hour rhythmic periodicity despite environmental temperature changes. While the molecular clockwork that drives genome-wide oscillations in gene expression is well characterized, how circadian gene expression contributes to the maintenance of rhythmic outputs under thermal variation and to produce temperature compensation remains uninvestigated. We profiled circadian gene expression in the ectothermic animal *Drosophila melanogaster* using time-series RNA-seq of head tissues under constant darkness at 18°C, 25°C, and 29°C. Locomotion assays confirmed that behavioral rhythms were temperature compensated under these conditions. Analyses of the time-series RNA-seq samples revealed hundreds of oscillating genes at each temperature, yet the majority were temperature specific. Only 14 genes, including the core clock components *period* (*per*) and *vrille* (*vri*), oscillated across all three constant darkness conditions at different temperatures. Even among these, the phase, amplitude, and expression levels of these oscillating genes often shifted with temperature. Thus, temperature-compensated oscillating genes are rare and largely confined to core clock genes, while most oscillating genes are temperature-specific. Our results suggested a potential architecture: a robust, temperature-compensated core set of oscillating genes, and a flexible, temperature-responsive output layer. The latter likely establishes temperature-specific oscillating gene expression programs that enable the animal to achieve temperature compensation at behavioral and physiological levels.

## Introduction

Circadian rhythms are ∼24-hour oscillations in metabolism, physiology, and behavior observed across animals, plants, fungi, and some prokaryotes. These rhythms enable organisms to adapt their life strategies to periodically fluctuating environments. Circadian phenomena share three defining properties. First, circadian oscillations are self-sustaining and persist under constant conditions. Second, the oscillations can be synchronized by environmental cues such as light and temperature. Third, unlike the temperature-sensitive rates of most biochemical reactions, the ∼24-hour period of circadian oscillations remains remarkably stable over a wide range of ambient temperatures, a phenomenon known as temperature compensation.

Temperature compensation ensures the reliability of the circadian clock, and this phenomenon has been recognized in bees, crabs, and clams even before the discovery of the circadian clock. Among these findings, Pittendrigh [1] experimentally demonstrated that the *Drosophila* eclosion rhythm retained a 24-hour period in constant darkness over a temperature range of 16°C-26°C. At the molecular level, the ATPase activity of KaiC in the unicellular photosynthetic cyanobacterium is also temperature compensated [2]. Most recently, circadian rhythms were identified in the nonphotosynthetic bacterium *Bacillus subtilis*, and these rhythms likewise display temperature compensation [3]. Moreover, circadian clocks in cell culture and tissue explants from homeothermic animals also exhibit temperature compensation [4-6]. These studies suggest temperature compensation as an evolutionarily adaptive mechanism for timekeeping, highlighting its fundamental importance in ensuring the clock’s stability across diverse organisms and biological contexts [7].

The molecular basis of the circadian clock has been extensively studied in model systems over the past four decades, including *Drosophila melanogaster*, *Mus musculus*, *Neurospora crassa*, and *Synechococcus elongatus* [8]. Forward mutagenesis screens in *Drosophila* led to the discovery and elucidation of the molecular clock, the core molecular components underlying the circadian clock [9]. Among these components, CLK (Clock) and CYC (Cycle) form a nuclear heterodimer that binds to the promoters of the *period*

*(per)* and *timeless (tim)* genes to activate their transcription. The resulting *per* and *tim* mRNAs are translated into proteins, PER (Period) and TIM (Timeless), in the cytoplasm. PER and TIM accumulate, dimerize, and translocate into the nucleus to inhibit CLK-CYC activity, thereby repressing their *(per and tim)* transcription. This transcriptional and translational feedback loop (TTFL) results in cyclic PER and TIM synthesis and degradation at ∼24-hour intervals, driving self-sustaining circadian rhythms. Light resets the clock via CRY (Cryptochrome), linking circadian rhythms to environmental photic cues [10]. Downstream of these feedback loops, the oscillatory expression of thousands of genes is regulated, orchestrating the clock’s output in various physiological and behavioral rhythms.

Despite the progress made in understanding the molecular clock, the mechanisms underlying circadian temperature compensation remain a mystery. Before the discovery of the molecular clock, a “network model” was proposed, suggesting that temperature-sensitive factors within the clock counteract each other to maintain a constant period under varying temperatures [11,12]. Another theoretical model, termed the “pathway model,” provided experimental evidence in *Drosophila* to support the idea that all period-determining components of the clock are independently temperature compensated. In this model, the core clock relies on specific pathways to convey temperature information, enabling individual components to scale properly [13,14]. While these models provided theoretical frameworks, experiments conducted using various clock mutants with defective rhythms suggested that protein-protein interactions, phosphorylation, nuclear translocation, and specific protein domains of the clock gene contribute to the stabilization of the TTFL and to temperature compensation [15-20].

Recent advances in genome-wide high-throughput analysis have enabled a comprehensive analysis of gene expression, revealing that the molecular clock not only regulates its own transcription, but also orchestrates genome-wide oscillations in gene expression, influencing diverse biological processes. For instance, nearly half of the protein-coding genes and over a thousand non-coding RNAs exhibit circadian transcription in *M. musculus* [21]. In *D. melanogaster* head tissues, hundreds of genes show circadian expression under the control of the core clock [22-24]. Genome-wide profiling of rhythmic gene expression has enabled a comprehensive investigation of the molecular clock’s impact on various downstream biological processes, thus offering new avenues for studying its output [25].

Such gene expression profiling has yet to be leveraged for studying temperature compensation. In this study, we examined whether temperature compensation applies to rhythmic expression of core clock genes and their output genes or pathways. Here, using time series RNA sequencing, we examined gene expression profiles in *Drosophila* heads every four hours over a total of two days under four conditions: light-dark cycles at 25°C, and constant darkness at 25°C, 18°C, and 29°C. The period of locomotor rhythm remained constant (∼24 hours), and temperature compensated across these conditions. We identified distinct sets of oscillatory genes specific to each temperature under constant darkness, each enriched for different gene ontology terms and pathways. This observation suggests that the downstream pathways of the molecular clock vary with temperature, and temperature compensation does not simply apply to all individual oscillating genes. In addition, we identified a small group of genes, consisting of core molecular clock genes and genes with unknown functions, that exhibited oscillatory expression under constant dark conditions across all tested temperatures, although their phase, amplitude, or overall expression level varied with temperature. These findings, derived from a large transcriptomics dataset, suggest a potential two-tier architecture: a robust, temperature-compensated core set of oscillating genes, and a flexible, temperature-responsive output layer. The latter likely establishes temperature-specific oscillating gene expression programs and pathways that enable the animal to drive 24-hour output rhythms at different temperatures.

## Materials and Methods

### Locomotor activity assay

*Drosophila melanogaster*, wild-type *Canton-S* strain, was used throughout the study. Flies were reared on standard molasses cornmeal and agar media with yeast at 25°C and under a 12-hour light/12-hour dark (12L:12D) condition. Male and female flies, aged 3 to 6 days, were briefly anesthetized with CO_2_ and placed into 5 x 65 mm glass tubes containing 5% sucrose and 2% agar. Locomotor activities of flies were monitored using the Drosophila Activity Monitor 2 (DAM2, Trikinetics, Waltham, MA, United States) under the following conditions: 3 days of 12L:12D at 25°C for entrainment, followed by 10 days of constant darkness (DD) at 18°C, 25°C, or 29°C for free-running. The visualization and calculation of actograms and periodograms were performed using the Rethomics R packages [26]. The periodogram properties, including power, period (in hours), and significance level, were calculated using the chi-square method.

### Sample collection

Flies were reared in bottles containing 50 mL of fly food under a 12L:12D cycle at 25°C. 20-30 male and 20-30 female flies were kept in each bottle. After 3 days of egg laying, the parental flies were removed from the bottles. Newly eclosed flies were collected and kept for 2 days under a 12L:12D at 25°C, followed by 3 days under 12L:12D at 25°C, DD at 18°C, DD at 25°C, or DD at 29°C before sample collections for the time series RNA-seq. Sample collections were performed every 4 hours over a 2-day period (48 hours), starting from ZT/CT0 to ZT/CT44. At each time point, male and female flies (approximately 200-300 flies per bottle) were collected, flash-frozen using liquid nitrogen, and stored at -80°C until RNA extraction. The 48hr time-course sample collections were performed twice, resulting in a total of 24 samples per condition per 2-day period and a total of 4 biological replicates per time-point per 24hr LD/DD cycle.

When collecting the samples, we used incubators with opposite light:dark lighting conditions. A 12L:12D incubator was used for entraining ZT/CT0, ZT/CT4, ZT/CT8, ZT/CT24, ZT/CT28, and ZT/CT32 samples, while a 12D:12L cycle incubator was used for entraining ZT/CT12, ZT/CT16, ZT/CT20, ZT/CT36, ZT/CT40, and ZT/CT44 samples.

### RNA sequencing

The fly samples were vigorously shaken to separate the fly heads from the bodies. The collected fly heads were then homogenized using a BeadBug™ 3 Microtube Homogenizer and prefilled 2.0 mL tubes filled with zirconium beads (Benchmark Scientific). Following the manufacturer’s protocols, total RNA was purified using the RNeasy Mini Kit (Qiagen), and genomic DNA was removed with RNase-free DNase (Qiagen). The cDNA library construction, including poly-A mRNA isolation, fragmentation, reverse transcription, and PCR amplification, was performed by Novogene Corporation Inc. using the NEBNext® Ultra™ II Directional RNA Library Prep Kit for Illumina® (New England Biolabs). The libraries were sequenced on the Illumina Novaseq 6000 sequencer to generate directional 150-bp paired-end reads.

### RNA sequencing data processing

Raw reads were filtered and trimmed using fastp (0.23.2) [27] with default parameters and subsequently aligned to the FlyBase reference genome (release r6.55) using HISAT2 (v2.2.1) [28]. Gene-level read counts were quantified using featureCounts [29] in the Subread package (v2.0.8). The quality control and mapping statistics of all RNA-Seq samples are summarized in Table S1. Read counts were normalized using DESeq2 (1.46.0) [30] to account for differences in sequencing depth and library size. Differentially expressed genes between conditions were identified using a generalized linear model implemented in edgeR (4.4.2) [31] (Table S3-5).

### Rhythmic gene expression analysis

Cycling transcripts were identified using JTK_CYCLE (v3.1) [32] with DESeq2 normalized counts. The analysis was conducted with a periodicity range of 20 to 28 hours, using 4-hour intervals over a 48-hour period, across two biological replicates for each of the four conditions. Genes with an adjusted *p*-value of < 0.05 were classified as oscillating genes (Table S2). The period length, amplitude, and peak timing of each oscillating gene were determined. Rhythmic features of shared oscillating genes were compared between conditions. Period and phase (peak timing) were directly compared, while amplitude differences were assessed using intermediate statistics (tmp) from JTK_CYCLE, which were used to estimate amplitude and evaluated with a median test. Other circadian rhythm detection tools, including eJTK_CYCLE [33], BiO_CYCLE [34], RAIN [35], and Lomb-Scargle (LS) [36], were used to validate the results obtained from JTK_CYCLE (Fig. S3).

### Functional enrichment analysis

Functional enrichment analysis was conducted using the online tool Metascape (http://metascape.org). Oscillating gene sets identified by JTK_CYCLE were analyzed across multiple functional sets, including Gene Ontology (GO), KEGG Pathway, and Reactome Gene Sets (Table S9). Terms meeting the criteria of a *p*-value < 0.01, a minimum count of 3, and an enrichment factor > 1.5 (where the enrichment factor represents the ratio of observed to expected counts) were identified. These terms were subsequently grouped into clusters based on their membership similarities to highlight functional relationships in gene enrichment heatmaps and dot plots.

## Results

### Rhythmic activity of *Drosophila melanogaster* is temperature compensated

To confirm the circadian rhythm of *Drosophila* locomotor activity is temperature compensated, the locomotor activities of wild-type *Canton-S* male and female flies were monitored using the Drosophila Activity Monitor (DAM) system (Fig.1A, C, and D and Fig. S2A, B, and C). The flies were initially monitored under a 12-hour light/12-hour dark (LD) cycle for two days. Subsequently, the conditions were changed to constant darkness (DD) conditions at 18°C, 25°C, or 29°C to assess the free-running periodicity under different temperature conditions, while a continuous 25°C LD cycle served as the control. For the first two days under the 25°C LD conditions (denoted as the entrainment phase in Fig. 1A), activity peaks were observed twice during daytime, at dawn and dusk. In contrast, under DD conditions, the activity pattern shifted, displaying a single peak within a 24-hour period, indicating that locomotor activity patterns were altered under DD conditions (denoted as the acclimation and sampling phases in Fig. 1A). To determine whether the period length (*tau*) of locomotor activity was maintained across different temperatures, activity data from six days under DD conditions were used to calculate *tau* using the chi-square periodogram [26] (Fig. 1C and D for males and Fig. S2B and C for females). The analysis revealed that a ∼24-hour free-running periodicity was consistently maintained across different temperature conditions, indicating that both male and female flies exhibit temperature compensation in their locomotor activity rhythm.

**Figure 1.**
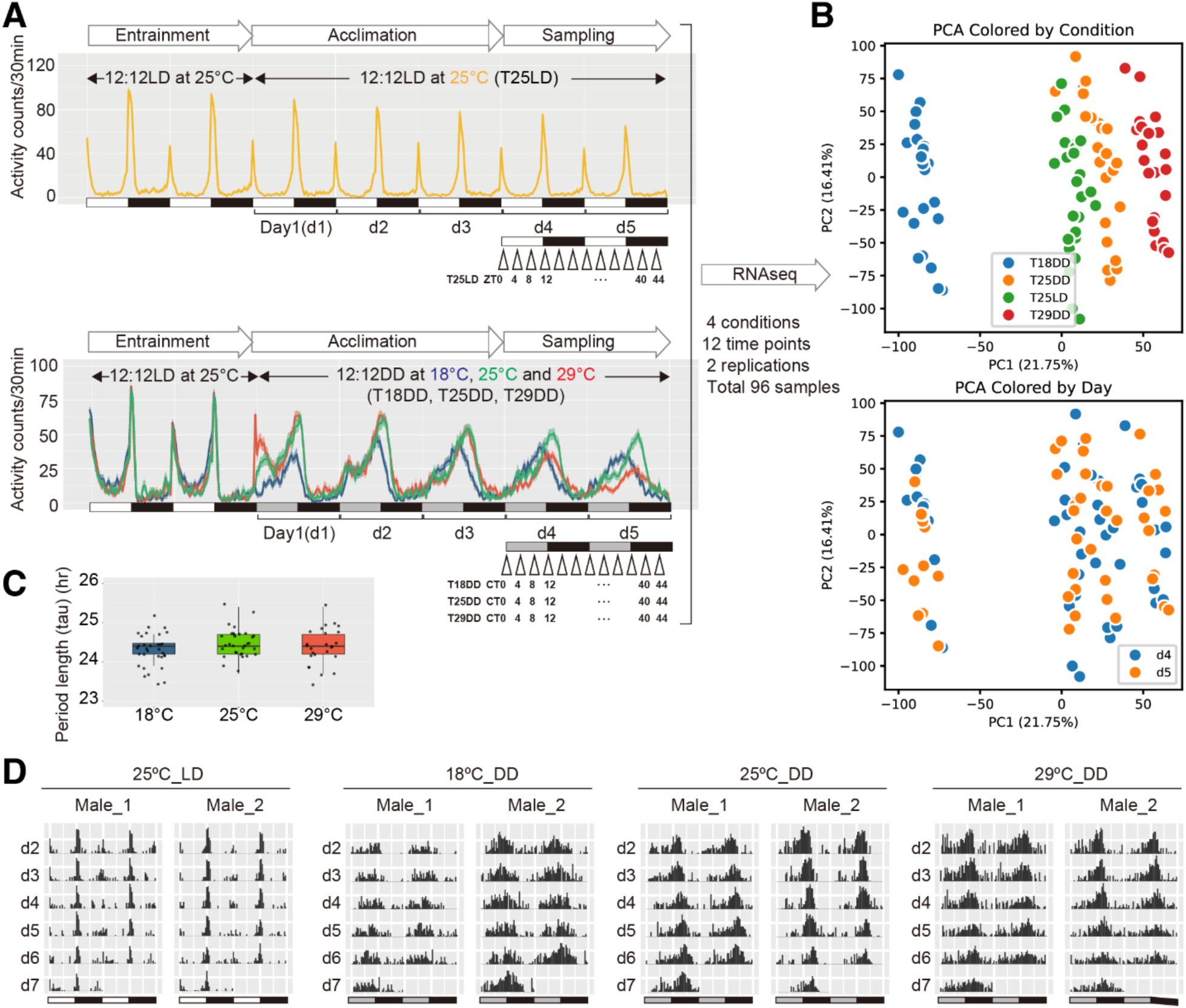
Overview of experiment design for time series RNA-seq sample collections under different temperature conditions. **(A)** Locomotor activity of male wild-type flies under the 25°C 12:12 LD condition (upper panel: T25DD, yellow) and three 12:12 DD temperature conditions (lower panel: T18DD, blue; T25DD, green; T29DD, red). Under the T25LD condition (upper panel), flies were monitored at 25 °C LD for two LD cycles (entrainment) before continuing under 25°C LD (acclimation) for three LD cycles (Day 1-Day 3, d1-d3). Fly head samples were collected every 4 h during the following two LD cycles (Day 4-Day 5, d4-d5), yielding 12 samples named T25LD ZT0–44. Under constant darkness (lower panel), flies were monitored at 25°C LD for two LD cycles (entrainment) before being transferred to 18°C DD, 25°C DD, or 29°C DD (acclimation) for three DD cycles (d1-d3). Fly head samples were collected every 4 h during the following two DD cycles (d4-d5), yielding 36 samples named T18DD CT0-44, T25DD CT0-44, and T29DD CT0-44. Sample collections for T25LD, T18DD, T25DD, and T29DD were repeated in an independent experiment. Across 4 conditions, 12 time points, and 2 replicates, a total of 96 samples were obtained to construct the time-series RNA-seq dataset. **(B)** PCA plots of the 96 RNA-seq samples, colored by collection condition (upper panel) or collection day (lower panel). **(C)** Scatter plots of estimated circadian period length (*tau*) of locomotor activity rhythm under the three DD conditions (Kruskal-Wallis test, *χ*² = 2.7145, *p* = 0.2574). **(D)** Representative double plots of locomotor activity of male flies under 25°C LD, 18°C DD, 25°C DD, and 29°C DD conditions.

### Time-series RNA Sequencing

Because oscillating genes, which are expressed in rhythmic patterns, are considered the output of the oscillating molecular clock, we next examined whether each of these oscillating genes is temperature compensated. To profile gene expression under DD conditions at different temperatures, time-series RNA sequencing was conducted using fly head samples. Wild-type *Canton-S* flies were grown under a 25°C LD condition. 2- to 4-day-old male and female flies were entrained for two days under a 25°C LD condition. Subsequently, they were moved to DD conditions at 18°C, 25°C, or 29°C for three days to allow acclimation. The first day under the DD condition was designated as Day1 (d1). After acclimation (Day1 to Day3, d1-d3), the fly heads were collected every four hours over a two-day period (Day4 and Day5, d4 and d5), with two biological replicates. Flies maintained under a continuous 25°C LD cycle condition were also collected on d4 and d5 and served as the control (Fig. 1A). In total, 96 samples were collected to analyze the rhythmic RNA expression patterns under temperature compensation (Fig. 1A, B, Fig. S2A, S2D, Fig. S1, and Table S1). Principal component analysis (PCA) was conducted to assess the samples (Fig. 1B and Fig. S2D). The PCA results demonstrated clustered patterns. Samples collected under the same lighting/temperature conditions clustered, and the clusters were separated from each other. In each cluster, samples from d4 and d5 intermingled. As expected, HEAT SHOCK PROTEIN (HSP) genes showed differential expression levels under different temperature conditions (Fig. S3 and Table S3-5). Most of the heat-induced HSP genes showed reduced expression when the 18°C DD condition was compared to the 25°C DD condition, while most of the heat-induced HSP genes showed increased expression when the 29°C DD condition was compared to the 25°C DD condition (Fig. S3A and B). Interestingly, a small number of HSP genes also changed their expression level when the 25°C LD condition was compared to the 25°C DD condition (Fig. S3A).

### Oscillating gene expression patterns under LD and DD conditions

Using the JTK_CYCLE algorithm [32], oscillating genes were identified under four conditions: 25°C LD, 25°C DD, 18°C DD, and 29°C DD. 1,733 genes exhibited rhythmic expression patterns under the 25°C LD condition. Under DD conditions, 291 genes oscillated at 18°C, 234 at 25°C, and 385 at 29°C, respectively (Fig. 2A and Table S2). Oscillating properties, including amplitude, *tau*, and phase, were summarized in Fig. 2B-D. After entering DD conditions for four days, oscillating genes exhibited reduced amplitude when the light cue was absent (Fig. 2B). The proportions of genes with a 20-hour, 24-hour, or 28-hour *tau* estimated by JTK_CYCLE were shown in Fig. 2C. Oscillating genes exhibited different phase distributions among different DD conditions (Fig. 2D). These results suggest that oscillating features of genes were influenced by temperature under DD conditions. While 1,517 oscillating genes were unique to the 25°C LD condition, 191, 126, and 273 were unique to the 18°C, 25°C, and 29°C DD conditions, respectively. The number of genes unique to two, three, or all four of the conditions was few (from 1 to 62, Fig. 2E).

**Figure 2.**
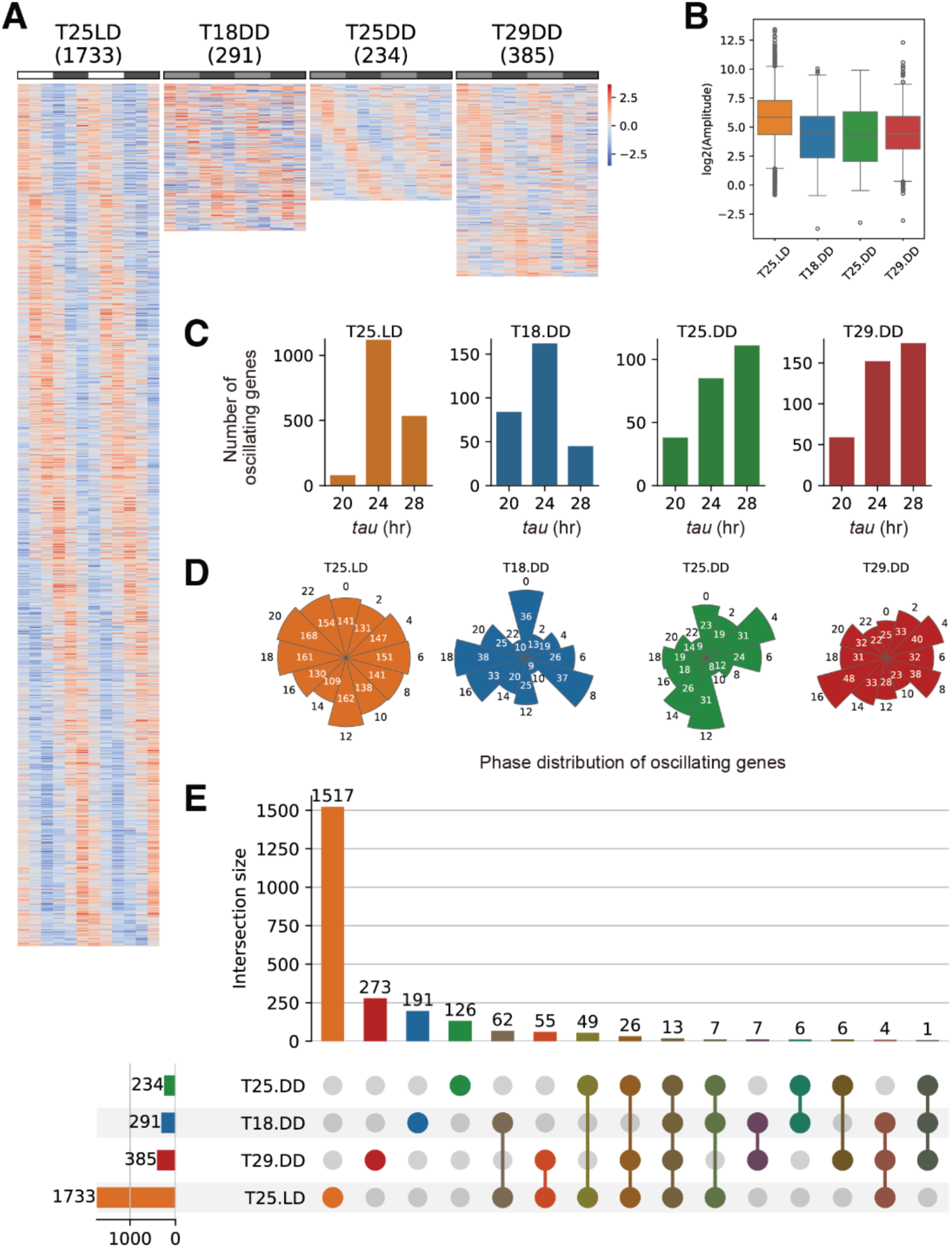
Identification of oscillating genes under different temperature conditions. **(A)** Heatmaps of the 1733, 291, 234, and 385 oscillating genes under 25°C LD,18°C DD, 25°C DD, and 29°C DD conditions, identified by JTK_CYCLE. **(B-D)** Amplitude (B), number (C), and phase distribution (D) of oscillating genes under 25°C LD,18°C DD, 25°C DD, and 29°C DD conditions. **(E)** Upset plots demonstrating the numbers of genes that were identified as oscillating genes under one, two, three, or all four conditions.

### Oscillating genes under different constant darkness temperature conditions

We next focused on analyzing rhythmic gene expression under DD conditions and found that 253, 175, and 328 genes exhibited temperature-specific oscillating patterns at 18°C, 25°C, and 29°C, respectively. These three groups of oscillating genes are temperature specific: they oscillated under one of the DD conditions but lost their oscillations completely under the other two temperature conditions. In contrast, only a small portion of the oscillating genes exhibited shared oscillating patterns: 32 genes were oscillating at both 29°C DD and 25°C DD; 13 genes were oscillating at both 18°C DD and 25°C DD; 11 genes were oscillating at both 18°C DD and 29°C DD. Only 14 genes were oscillating across all three DD conditions (Fig. 3A, C, and Table S2).

**Figure 3.**
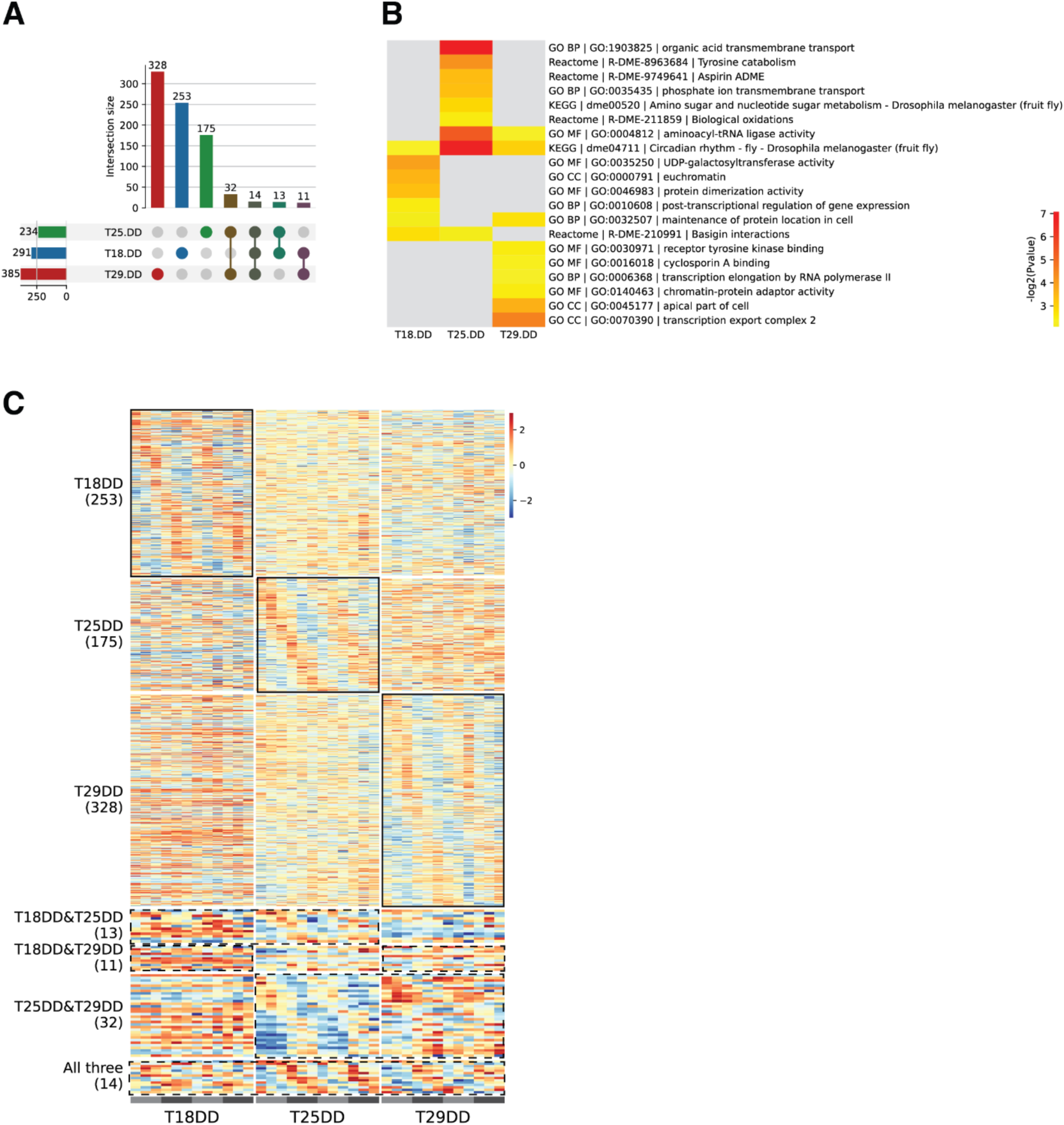
Identification of temperature-specific and temperature-compensated oscillating genes. **(A)** Upset plots demonstrating the numbers of genes that were identified as oscillating genes under one, two, or all three constant darkness conditions (18°C DD, 25°C DD, and 29°C DD). **(B)** Gene Ontology (GO) enrichment analysis of oscillating genes under 18°C DD, 25°C DD, and 29°C DD conditions. **(C)** Heatmaps of oscillating genes unique to each of the 18°C DD, 25°C DD, and 29°C DD conditions. These genes were considered temperature-specific oscillating genes. Heatmaps of shared oscillating genes: genes that were identified as oscillating genes under at least two of the three constant darkness conditions (18°C DD & 25°C DD, 18°C DD & 29°C DD, 25°C DD & 29°C DD, and 18°C DD & 25°C DD & 29°C DD). The genes that preserve a ∼24-hour rhythm under the 18°C DD & 25°C DD & 29°C DD conditions were considered temperature-compensated genes.

To understand if rhythmically expressed genes under the three different DD conditions contribute to specific pathways, we conducted functional enrichment analyses. Consistent with the oscillating genes being temperature-dependent, oscillating genes under each temperature condition were categorized into distinct functional groups (Fig. 3B, Fig. S5, and Table S9). Functional groups enriched for one specific temperature were rarely enriched for the other two temperature conditions. Interestingly, the GO term related to “circadian rhythm” was the only term shared across all three temperature conditions, indicating that the 14 oscillating genes shared across all three DD conditions are enriched for “circadian rhythm.” This finding indicated that the 14 temperature-compensated genes are likely those driving the circadian rhythm rather than output genes. Time-series RNA sequencing under the three DD temperature conditions, therefore, revealed that while the behavioral rhythmicity is maintained across different temperature conditions, the underlying oscillating gene groups and their regulatory pathways are largely temperature dependent or temperature specific.

### Temperature-compensated oscillating genes

Temperature compensation specifically refers to the periodicity of the circadian oscillation, which remains stable over a wide range of ambient temperatures. We reasoned that, to qualify as a temperature-compensated gene, a gene must oscillate at all three DD conditions. Accordingly, we define temperature-compensated genes as those that oscillated at 18°C DD, 25°C DD, and 29°C DD conditions (Fig. 3A, C, and Fig. 4). Strictly speaking, only genes that oscillate with the same *tau* under all temperature conditions can be considered truly temperature compensated, but such a stringent requirement would yield even fewer candidates. We therefore focused our discussion on these 14 genes.

**Figure 4.**
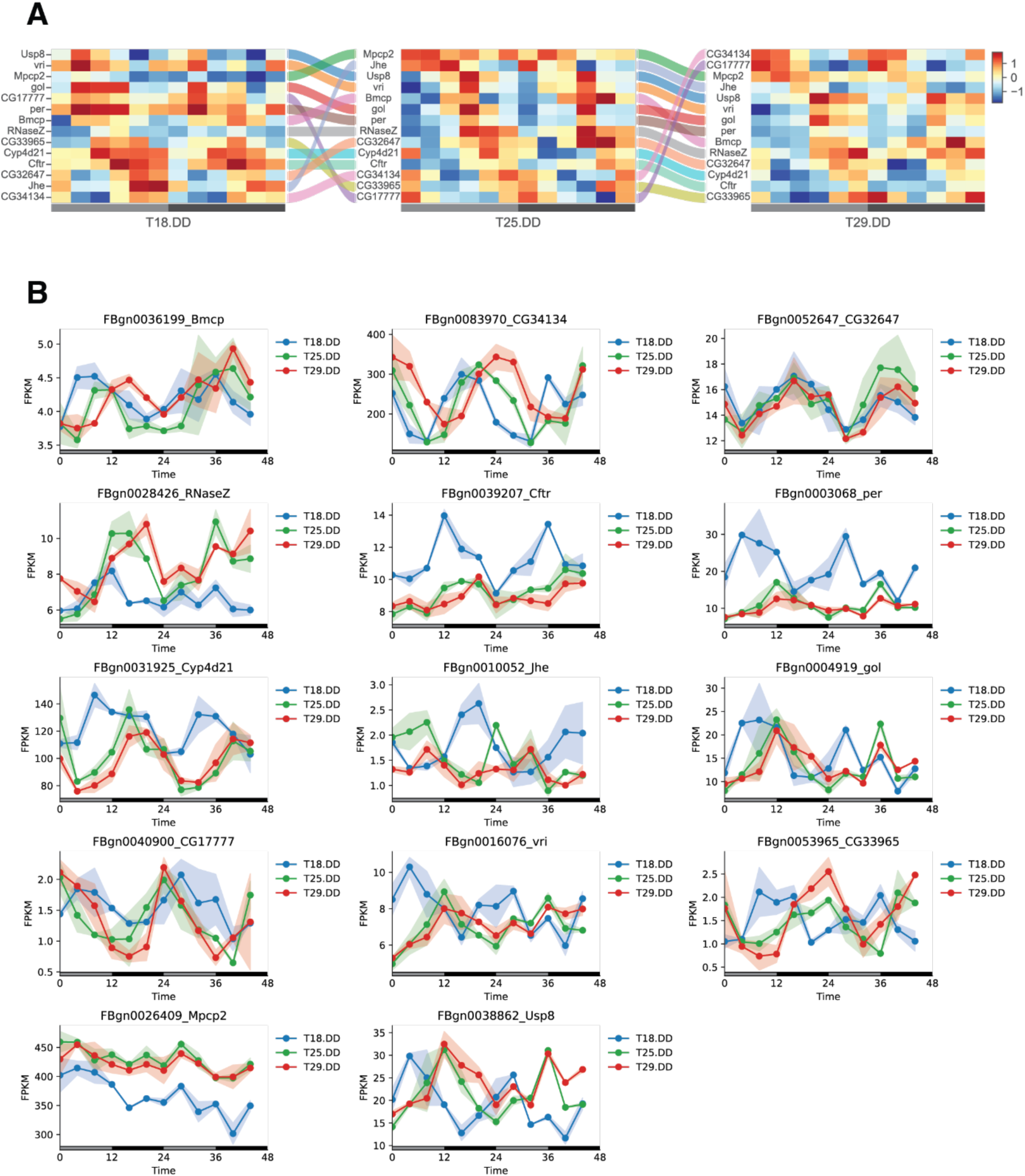
Expression patterns of 14 temperature-compensated genes. **(A)** Sankey diagram showing how the oscillatory properties of each of the 14 temperature-compensated genes change in relation to one another. In each heatmap for a given temperature condition, genes are ordered by phase, and identical genes are linked across conditions by flow lines. **(B)** Expression patterns of the 14 temperature-compensated oscillating genes across three temperature conditions. Lines represent mean expression levels.

When visualized using a Sankey diagram to display these genes based on the phase of their oscillations at each temperature condition, the data show that each of the 14 genes alters its ranking across the three heatmaps corresponding to the three temperature conditions (Fig. 4A). For example, gene *CG34134* oscillated at 18°C DD with a peak around CT16, at 25°C DD with a peak around CT20, and at 29°C DD with a peak around CT24 (CT0) (Fig. 4).

Among the 14 temperature-compensated genes, *period* (*per*) and *vrille* (*vri*) are well-characterized molecular clock genes [37-39]. Interestingly, both *per* and *vri* exhibited a phase advancement and increased expression levels at 18°C DD. In addition, *Cyp4d21*, *Usp8*, and *gol* have previously been identified as regulators of the circadian clock or clock-regulated genes [40-42]. The rest of the 14 genes consist of those with less-known functions but exhibited interesting oscillating patterns. For example, *CG32647* and *CG17777* exhibited stable oscillating patterns across all three conditions, while *CG34134* appeared to shift its peak timing as temperature changed. *CG34134* was previously reported as an oscillating gene in a study using dense time-course gene expression profiling to study *Drosophila* innate immune response [43]. An ABC transporter gene, *Cftr*, was also detected as temperature compensated across the three DD conditions. *Cftr* was recently predicted to encode the *Drosophila* equivalent of human CFTR, Cystic Fibrosis Transmembrane Conductance Regulator [44]. We did not observe strong or shared oscillating patterns in other known clock genes, such as *timeless*, *Clock, cryptochrome*, and *clockwork orange* [45-50]. This may reflect dampened rhythms in whole-head tissues after four days in constant darkness (DD). Specific isoforms of molecular clock genes might still oscillate strongly, as alternative splicing of clock genes has been shown to mediate temperature responses in *Drosophila* [40]. Although we could not assess splicing isoform-level gene oscillations, long-read RNA Sequencing may address this in the future.

To facilitate data exploration, we have made the time-series RNA-seq data available through an open-access, interactive web portal (https://drosophilarhythmicgenes.onrender.com). Users can easily examine the expression and oscillation patterns of any gene under different temperature conditions.

We next focused on the shared oscillating genes between any two of the three DD temperature conditions, and examined their period, phase, amplitude, and overall expression level (Fig. 5, Table S6-8). For example, among the 27 genes that exhibited oscillating gene expression at 18°C DD and 25°C DD, 14 maintained an unchanged periodicity, 16 exhibited different phases, and 3 exhibited significantly different amplitude (Fig. 5A). Similarly, among the 25 genes that exhibited oscillating gene expression at both 18°C DD and 29°C DD, 20 exhibited different phases. Among the 46 genes that exhibited oscillating gene expression at both 25°C DD and 29°C DD, 28 exhibited different phases. These results indicate that phase is particularly sensitive to temperature variations, a pattern further supported by the Sankey diagrams (Fig. S6).

**Figure 5.**
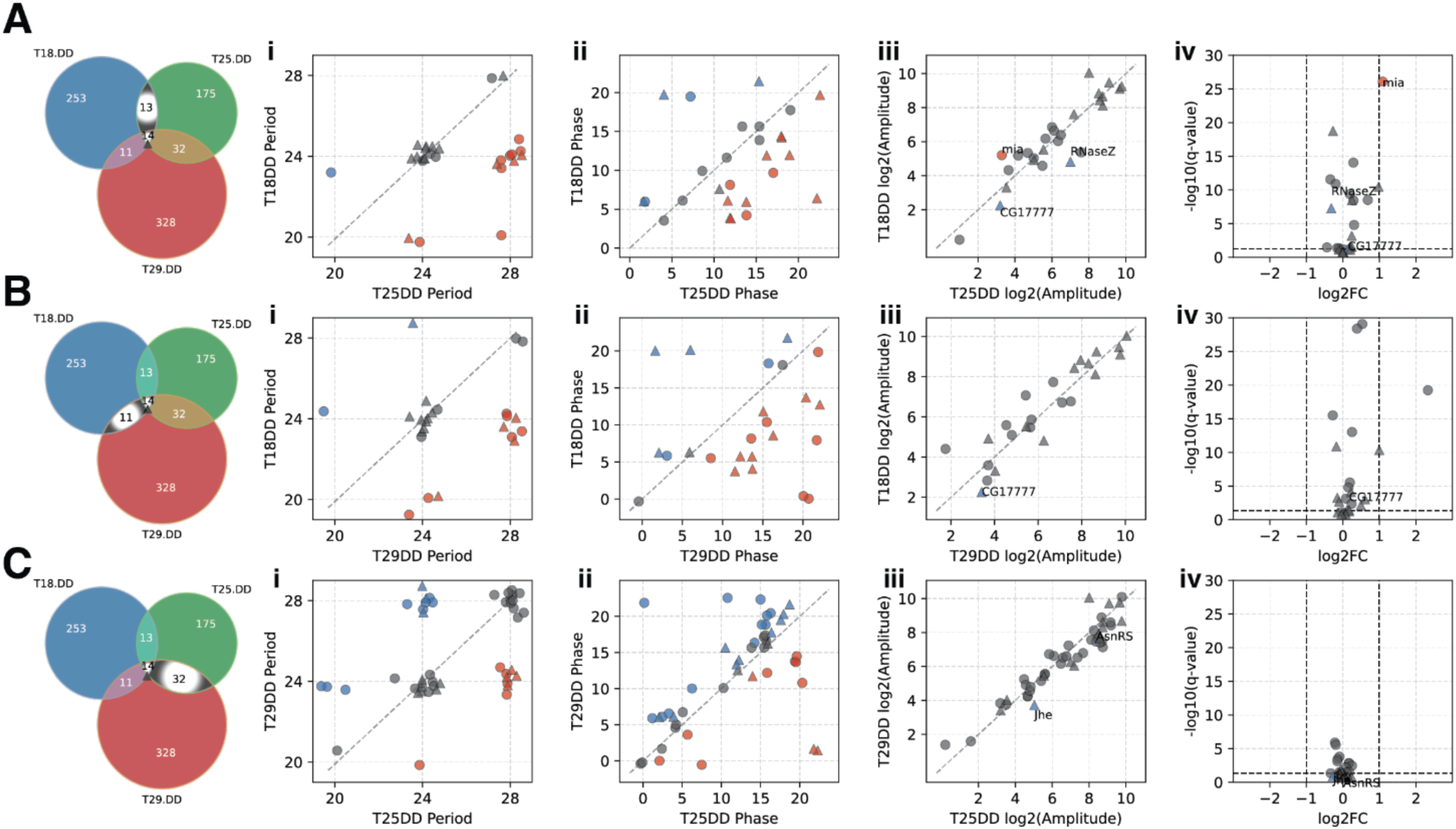
Comparison of oscillation properties between temperature conditions. **(A)** Comparisons of oscillation properties for the 27 oscillating genes shared between 18°C DD and 25°C DD. **(B)** Comparisons of oscillation properties for the 25 oscillating genes shared between 18°C DD and 29°C DD. **(C)** Comparisons of oscillation properties for the 35 oscillating genes shared between 25°C DD and 29°C DD. **(A-C, i-iv)** The properties being analyzed for these genes are: period length (i), phase (ii), amplitude (iii), and differential gene expression pattern (iv). In all four comparisons (i-iv), the triangles in the plots represent the 14 temperature-compensated oscillating genes, while the circles represent additional genes that oscillate in the two compared DD conditions but not in the third. The genes with changes are highlighted in different colors: red indicates a lengthened period (i), delayed phase (ii), and significantly increased amplitude (iii, *p*<0.05); blue indicates a shortened period (i), advanced phase, and significantly decreased amplitude (iii, *p*<0.05). Genes with significantly changed amplitude were also highlighted in the volcano plot of differential gene expression (iv).

## Discussion

Time-series RNA-seq has become a powerful method for studying circadian biology, enabling mechanistic insights into how environmental cues shape the circadian clock and its behavioral and physiological outputs. However, this approach has not been widely applied to the study of temperature compensation, a longstanding mystery in chronobiology. Previous studies have primarily focused on core clock genes and behavioral rhythms, showing that their ∼24-hour periodicity persists across a range of constant temperatures in darkness. In this study, we expanded the scope to include hundreds of oscillating genes whose rhythmic expression is thought to underlie circadian patterns of behavior and physiology. We examined whether these individual genes are temperature compensated. Surprisingly, only a limited subset retained oscillatory behavior across all three DD conditions, suggesting that temperature compensation is rarely preserved at the individual gene level. Although these genes maintained a ∼24-hour period, their phase, amplitude, or expression level often shifted with temperature. This small group of temperature-compensated genes comprises both well-characterized core clock components and genes not previously linked to circadian function. We propose that these genes may contribute to the core molecular oscillator, consistent with its well-established temperature compensation across diverse systems.

Our analysis also reveals distinct sets of genes with oscillating expression patterns unique to each temperature condition. These temperature-specific gene sets are enriched for different biological pathways, indicating that distinct molecular processes are engaged at different temperatures. Moreover, among the genes that oscillate under multiple temperature conditions, key oscillatory properties, particularly phase, shift in response to temperature changes. The hundreds of temperature-specific oscillating genes observed at each condition highlight how the circadian system recruits different transcriptional programs to adapt to environmental temperature. These findings suggest that ambient temperature can dynamically reshape the oscillatory transcriptome: existing rhythmic genes may lose their oscillation, while previously non-oscillating genes may become rhythmic as temperature fluctuates.

In summary, our study provides new insights into temperature compensation by applying time-series RNA-seq to circadian gene expression analysis. When first proposing the TTFL model, Hardin *et al.* (1990) noted that the presence of feedback regulation makes it difficult to clearly distinguish core clock function from its downstream outputs [51]. Our findings suggest that temperature compensation may serve as a useful criterion for making this distinction. Specifically, we find that a small set of temperature-compensated oscillating genes likely defines the core clock mechanism. These genes interact with temperature-sensing pathways, adjust their oscillatory properties, modulate distinct biological programs, and ultimately shape temperature-specific clock outputs. Our findings also suggest that circadian temperature compensation is achieved through temperature-dependent processes. Along these lines, recent work in cyanobacteria suggested that the temperature-dependent fold-switching mechanism of KaiB contributed to the temperature-compensated KaiC ATPase activity [52].

This core-output division offers a conceptual bridge between molecular clockwork and environmental adaptation. Future studies at higher spatial and temporal resolution may further illuminate the mechanisms of temperature compensation. For example, time-series RNA-seq from specific tissues or even single cells under different temperature conditions could distinguish temperature-compensated genes from temperature-specific ones across clock and non-clock cells. Additionally, continuous monitoring of individual gene expression could reveal overcompensated or undercompensated genes: those with circadian periods slightly shorter or longer than 24 hours (e.g., 23.6 or 24.5 hours).

## Data, Materials, and Software Availability

Raw RNA-seq reads and processed data files have been deposited to the Gene Expression Omnibus (GEO) under accession number GSE302579. The time-series RNA-seq data is also available through an open-access and interactive web portal (https://drosophilarhythmicgenes.onrender.com).

## Acknowledgements

This work was supported by the Cancer Prevention and Research Institute of Texas (RR220021 to W.L.) and the National Institute of General Medical Sciences (GM150832 to W.L.). Portions of this research were conducted with the advanced computing resources provided by Texas A&M High Performance Research Computing. We thank Paul Hardin, Christine Merlin, Yuchao Jiang, and Jerome Menet for advice on experiment design and bioinformatics analysis. We thank Xin Chen, Fiona Gugala, Doris Migliaccio, Sydney Christensen, and Min Feng for comments on the manuscripts. We thank members of the Li laboratory for technical support.

## Author Contributions

F.I. and W.L. conceived and designed research; F.I. performed time-series RNA-seq sample collections and behavior experiments/analyses; F.Z., F.I., and W.L. conducted bioinformatics and statistical analyses; W.L., F.I., and F.Z. wrote the manuscript.

## Tables

**Table S1. Summary of RNA-seq data quality and mapping statistics**

File name: TableS1_RNAseq_QC_Mapping.xlsx

**Table S2. Oscillating genes identified using JTK_CYCLE**

File name: TableS2_JTK_CYCLE.result.xlsx

**Table S3. Results of differential gene expression between T18.DD and T25.DD**

File name: TableS3_T18.DD_vs_T25.DD.diff.result.xlsx

**Table S4. Results of differential gene expression between T29.DD and T25.DD**

File name: TableS4_T29.DD_vs_T25.DD.diff.result.xlsx

**Table S5. Results of differential gene expression between T25.LD and T25.DD**

File name: TableS5_T25.LD_vs_T25.DD.diff.result.xlsx

**Table S6. Amplitude comparison of shared oscillating genes under T18.DD and T25.DD**

File name: TableS6_T18DD_vs_T25DD.amplitude_comparison_results.xlsx

**Table S7. Amplitude comparison of shared oscillating genes under T18.DD and T29.DD**

File name: TableS7_T18DD_vs_T29DD.amplitude_comparison_results.xlsx

**Table S8. Amplitude comparison of shared oscillating genes under T29.DD and T25.DD**

File name: TableS8_T29DD_vs_T25DD.amplitude_comparison_results.xlsx

**Table S9. Gene Ontology enrichment analysis of oscillating genes under DD conditions**

File name: TableS9_Enrichment_AllLists.xlsx

## Supplementary Figures

**Figure S1.**
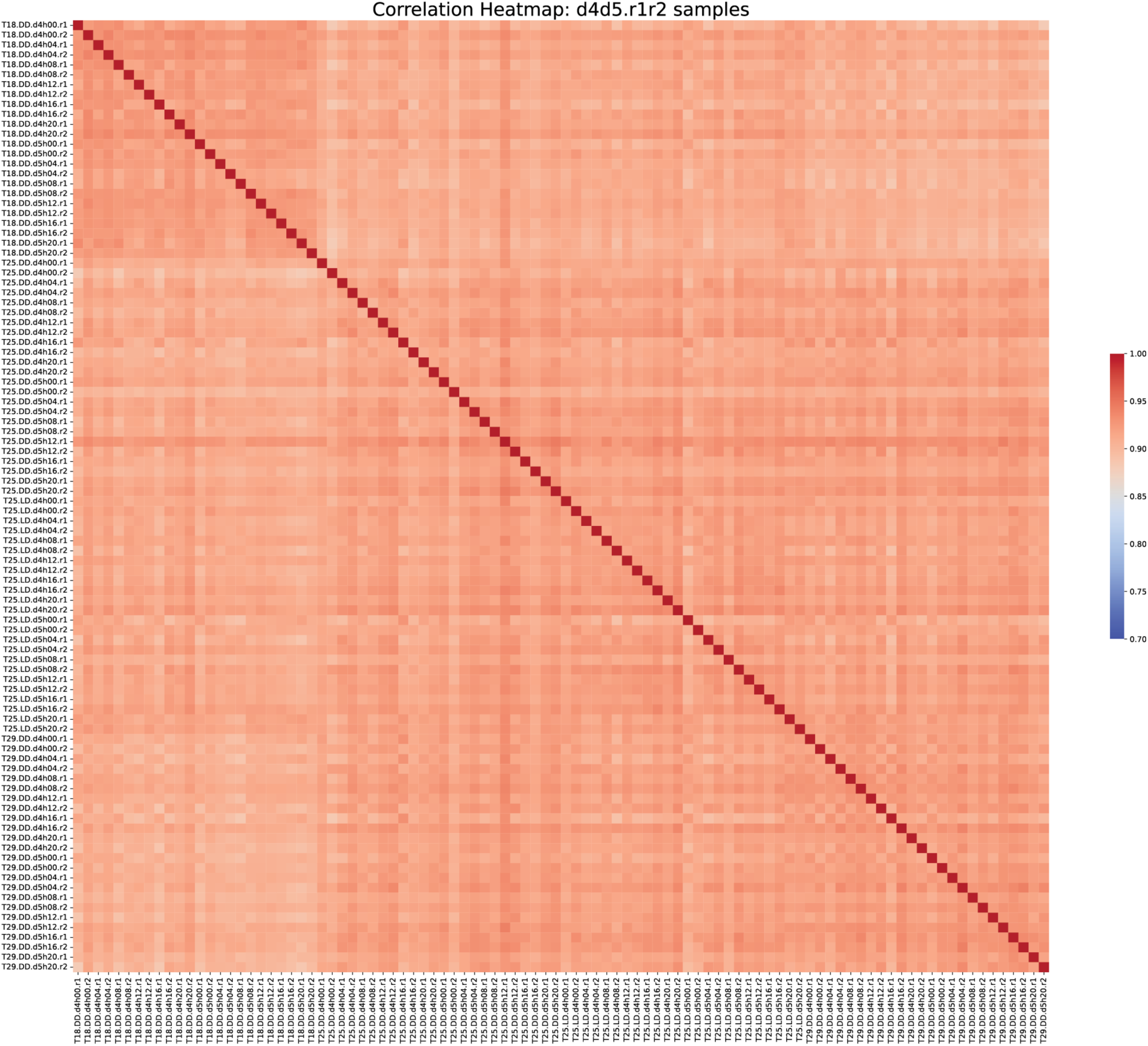
Correlation matrix of the 96 time-series RNA-seq samples.

**Figure S2.**
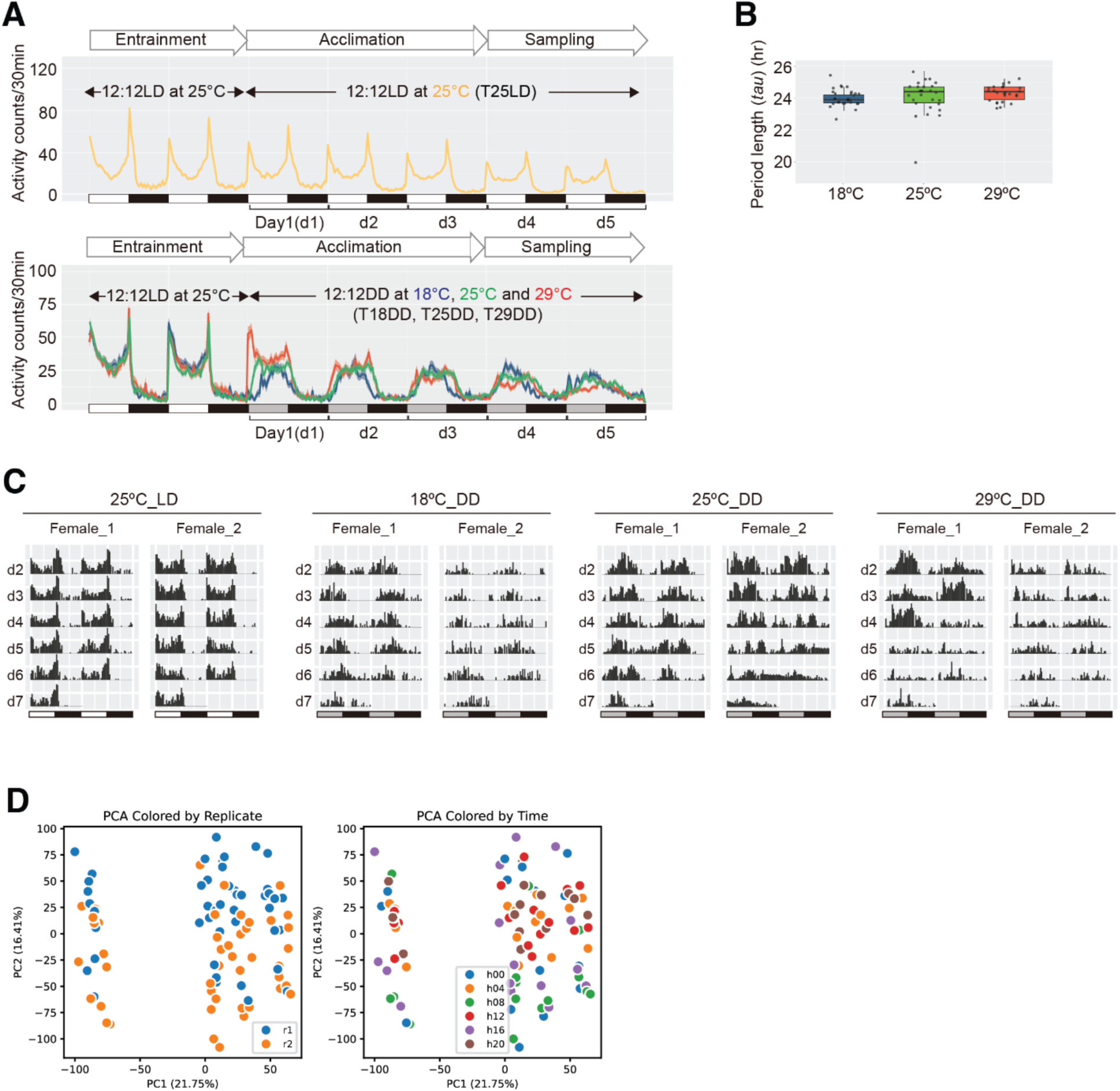
Overview of experiment design for time series RNA sequencing sample collections under different temperature conditions and locomotor activity of female animals under these conditions. **(A)** Locomotor activity of female wild-type flies under the 25 °C 12:12 LD condition (upper panel: T25DD, yellow) and three 12:12 DD temperature conditions (lower panel: T18DD, blue; T25DD, green; T29DD, red). Under the T25LD condition (upper panel), flies were monitored at 25 °C LD for two LD cycles (entrainment) before continuing under 25 °C LD (acclimation) for three LD cycles (Day 1–Day 3, d1–d3). Under constant darkness (lower panel), flies were monitored at 25 °C LD for two LD cycles (entrainment) before being transferred to 18 °C DD, 25 °C DD, or 29 °C DD (acclimation) for three DD cycles (d1–d3). **(B)** PCA plots of the 96 RNA-seq samples, colored by collection condition (upper panel) or collection day (lower panel). **(C)** Representative double plots of locomotor activity of female flies under 25°C LD, 18°C DD, 25°C DD, and 29°C DD conditions. **(D)** PCA plots of the 96 RNA-seq samples, colored by batch of replicate (left panel, replicate 1, r1; replicate 2, r2) or timepoint (right panel, h00 includes ZT00, ZT24, CT00, and CT24; h04 includes ZT04, ZT28, CT04, and CT28; h08 includes ZT08, ZT32, CT08, and CT32; h12 includes ZT12, ZT36, CT12, and CT36; h16 includes ZT16, ZT40, CT00, and CT40; h20 includes ZT20, ZT44, CT20, and CT44).

**Figure S3.**
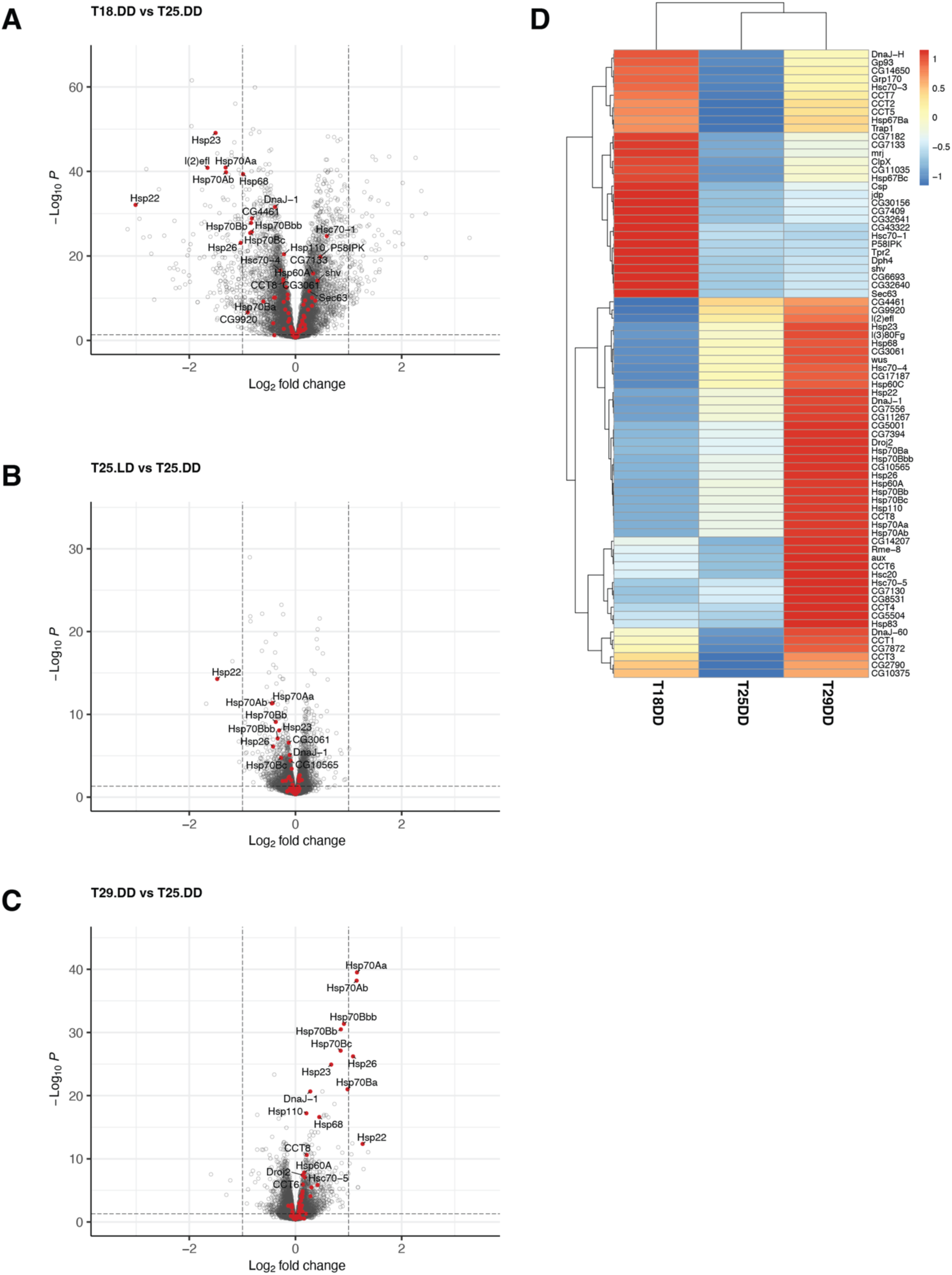
Differential gene expression analysis between constant darkness conditions. (A-C) Volcano plots of differential gene expression analysis. Comparison between 18 °C DD and 25 °C DD conditions (A), comparison between 25 °C LD and 25 °C DD conditions (B), and comparison between 29 °C LD and 25 °C DD conditions. Red dots indicate genes belonging to the gene group of HEAT SHOCK PROTEINS (HSP) (FlyBase ID: FBgg0000501). **(D)** Heatmap showing the expression levels of the HSP genes under the three constant darkness conditions.

**Figure S4.**
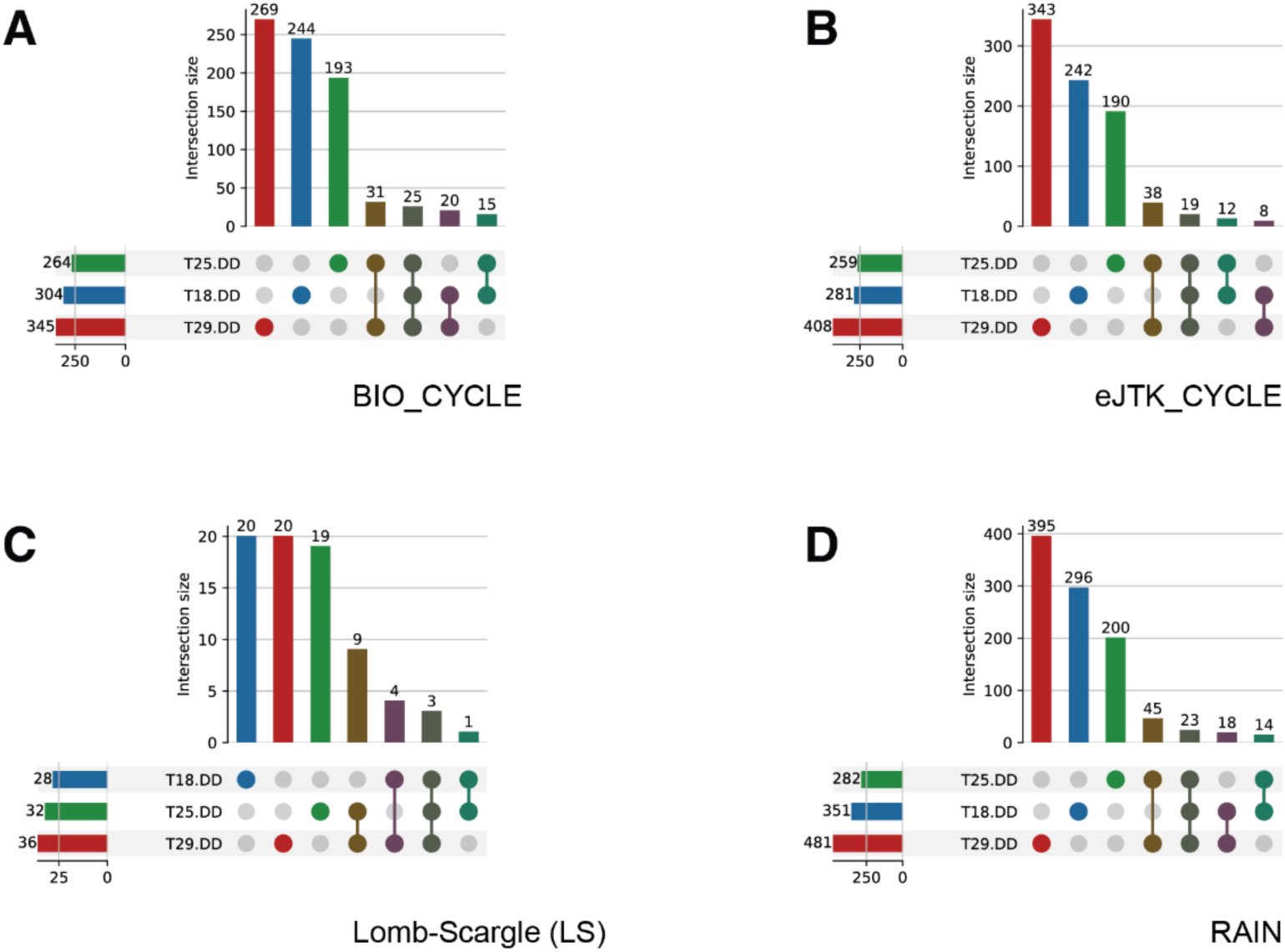
Numbers of oscillating genes identified using additional algorithms. (A-D) Upset plots showing numbers of genes that were identified as oscillating genes under one, two, or all three DD conditions, using BIO_CYCLE (A), eJTK_CYCLE (B), Lomb-Scargle (LS) (C) and RAIN (D).

**Figure S5.**
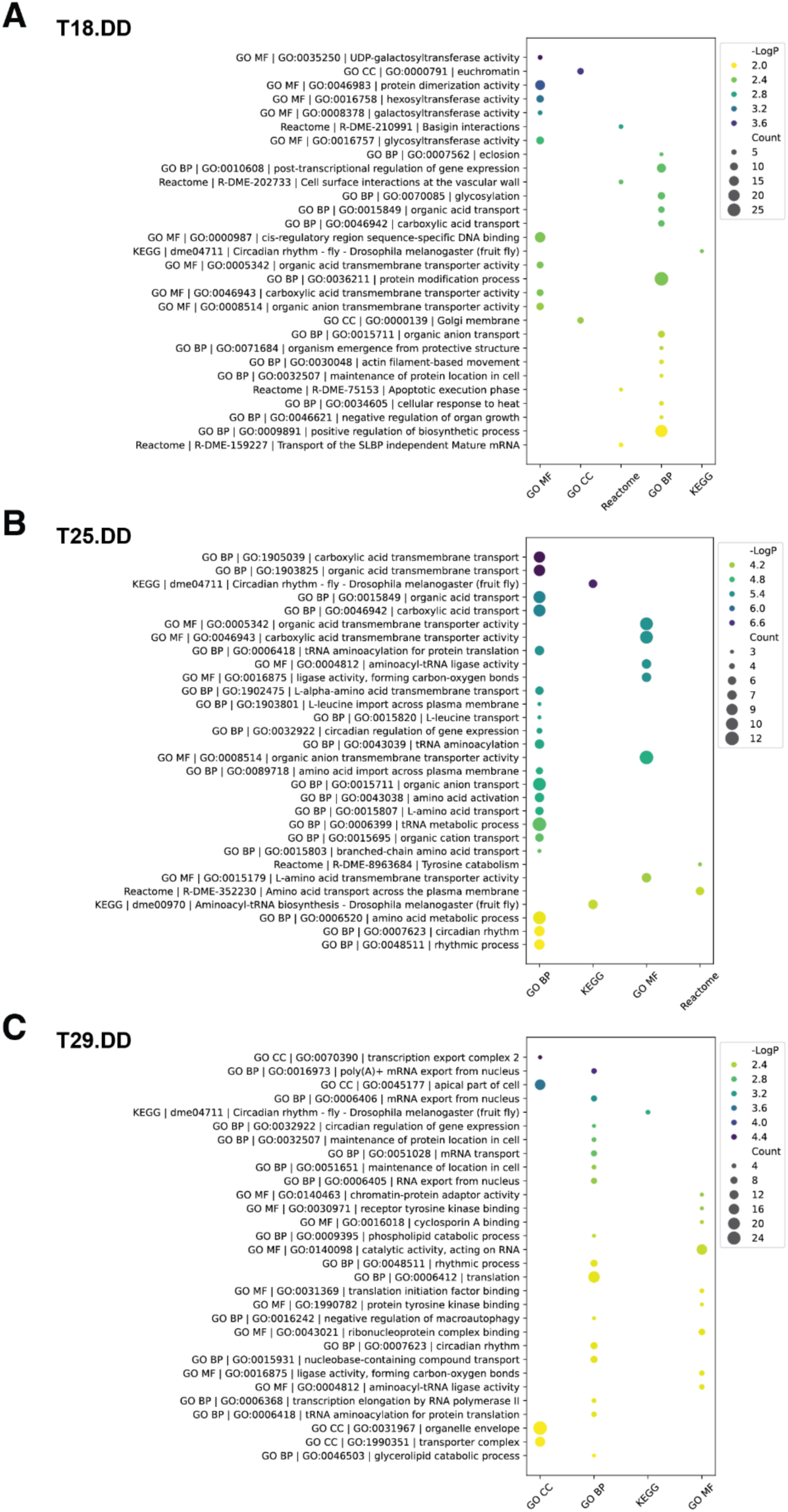
Gene ontology enrichment analysis for oscillating genes under each of the constant darkness conditions. **(A-C)** Oscillating genes under 18°C DD (A), 25°C DD (B), and 29°C DD (C) conditions were analyzed for enrichment using Gene Ontology (GO), KEGG Pathway, and Reactome Gene Sets. The identified enriched terms were plotted based on their *p*-value and count.

**Figure S6.**
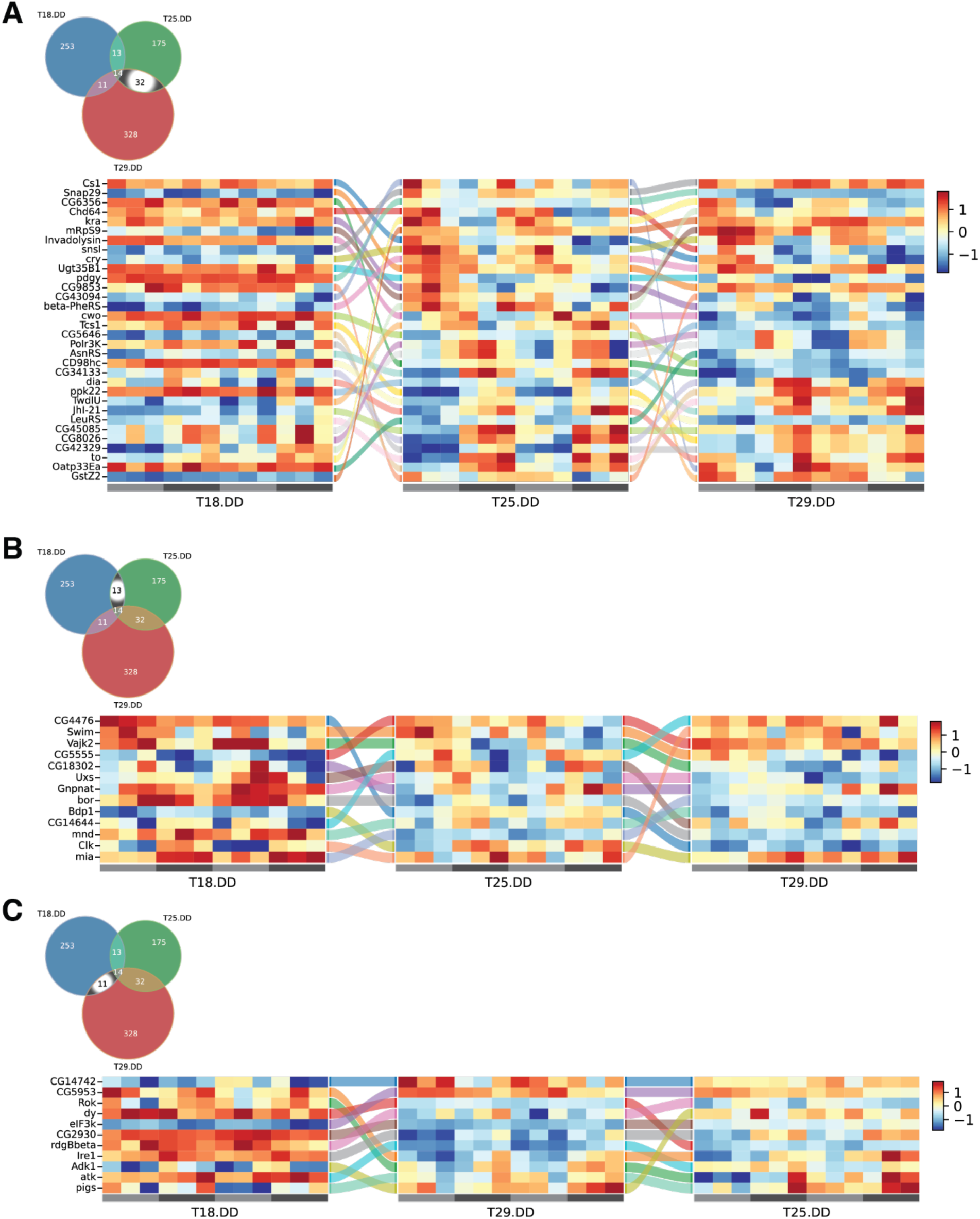
Shared oscillating genes under two of the three constant darkness conditions. **(A)** 32 genes were identified as shared oscillating genes under 25°C DD and 29°C DD conditions, but not under the 18°C DD condition. **(B)** 13 genes were identified as shared oscillating genes under 18°C DD and 25°C DD conditions, but not under the 29 °C DD condition. **(C)** 11 genes were identified as shared oscillating genes under 18°C DD and 29°C DD conditions, but not under the 25°C DD condition. **(A-C)** The Sankey diagrams illustrate how the oscillatory properties of these genes (A, 32 genes; B, 13 genes; C, 11 genes) change relative to each other. In each temperature-specific heatmap, genes are ordered by phase, and identical genes are connected across conditions by flow lines.

